# Non-Covalent Poly(ADP-ribose) Signaling Organizes a Circadian E3 Ligase Network in the Brain

**DOI:** 10.64898/2026.08.30.748055

**Authors:** Sajiv Harikrishnan, Sung-Ung Kang

**Affiliations:** Department of Biomedical Engineering, Johns Hopkins University, Baltimore, MD 21218, USA; Neuroregeneration and Stem Cell Programs, Institute for Cell Engineering, Johns Hopkins University School of Medicine, Baltimore, MD 21205, USA; Department of Neurology, Johns Hopkins University School of Medicine, Baltimore, MD 21205 USA

**Author notes:** Correspondence should be addressed to: Neuroregeneration and Stem Cell Programs Institute for Cell Engineering, Johns Hopkins University School of Medicine, 733 North Broadway, Suite 713, Baltimore, MD 21205.

**Keywords:** Non-covalent PAR signaling, PAR-binding proteins, E3 ubiquitin ligases, circadian biology, proteostasis, neurodegeneration, microglia

## Abstract

Although PAR biology has traditionally been studied through covalent PARylation, non-covalent PAR-binding proteins provide an additional mechanism for interpreting transient PAR signals and converting them into downstream regulatory programs. Among these effectors, E3 ubiquitin ligases are uniquely positioned to couple PAR sensing to selective ubiquitination, thereby integrating stress signaling with proteostatic control. Because circadian systems depend heavily on temporally coordinated protein turnover, we hypothesized that PAR-binding E3 ligases may form a circadian-structured regulatory layer within the brain. To test this, we integrated GTEx v10 brain transcriptomics, GWAS Catalog gene-mapped associations, CIRCA circadian phase annotations, and Human Protein Atlas single-cell transcriptomic resources to characterize the organization of PAR-binding E3 ubiquitin ligases across neural tissues. Across the brain, the E3 ligase repertoire was broadly deployed yet regionally structured, with cerebellar and cortical enrichment patterns preserved within the PAR-binding subset. Representative ligases spanning circadian regulation, DNA repair, and neurodegeneration-relevant pathways displayed distinct abundance and regional-variability archetypes across GTEx brain regions. Human genetic analyses demonstrated that E3 ligases associated with cognition-, neurodegeneration-, and sleep/circadian-related phenotypes were disproportionately PAR-binding, supporting convergence between PAR-responsive ubiquitin regulation and disease-relevant biology. Circadian phase analyses further revealed that PAR-binding ligases occupy structured, non-random circadian windows within the broader E3 background, including distinct co-phasing relationships with BMAL1 and CRY1. Finally, cell-type enrichment analyses identified microglia as the dominant compartment for circadian-linked and PAR-binding circadian E3 weighting within the brain E3 program. Together, these findings support a systems-level framework in which non-covalent PAR-binding E3 ubiquitin ligases constitute a brain-deployed, circadian-organized regulatory layer that couples PAR signaling to time-dependent ubiquitin control in neural systems.

## Introduction

Poly(ADP-ribose) (PAR) is a highly versatile signaling molecule that regulates processes ranging from the immediate response to DNA damage to longer-term control of chromatin structure, transcription, and nuclear organization.^1^ Synthesized primarily by poly(ADP-ribose) polymerase-1 (PARP1) in the nucleus, this negatively charged polymer acts as a potent signaling platform.^1^ It functions through either covalent PARylation of target proteins or non-covalent interactions that recruit and modulate PAR “reader” proteins as allosteric regulators.^2,3^ A broad array of proteins has evolved to interpret this PAR signal using specialized binding domains and physical motifs, forming a dynamic and context-dependent network often described as the PAR signalosome/interactome.^2,3^ Because PAR synthesis directly consumes cellular NAD⁺ and scales rapidly with stress intensity, PAR signaling is positioned not only as a DNA damage response component but also as an integrator of metabolic state, chromatin regulation, and cell-fate decisions.^1^

In parallel, the ubiquitin–proteasome system provides a complementary control layer that governs protein stability, localization, and signaling competence through targeted ubiquitination. E3 ubiquitin ligases are fundamental regulators of this process, acting as substrate-selective enzymes that couple substrate recognition to ubiquitin transfer by E2 conjugating enzymes. In neural tissues, where proteostatic integrity must be maintained over long time scales under substantial metabolic and oxidative burden, E3 ligases contribute to protein quality control, repair complex turnover, synaptic remodeling, mitochondrial homeostasis, and stress adaptation. While many E3 ligases function through canonical substrate recruitment and ubiquitin transfer, a subset incorporates non-covalent binding interfaces with PAR or PARylated assemblies.^3^ Compared to covalent PARylation, non-covalent PAR binding can act as a transient switch. It can recruit E3 ligases to PAR-rich compartments, allosterically modulate ligase activity, and/or bias substrate selection toward PARylated or PAR-associated proteins.^4^ This PAR–E3 coupling therefore provides a mechanistic route by which PAR signaling can be converted into selective ubiquitination programs that resolve stress responses, reshape chromatin-associated protein networks, and enforce proteostatic decision-making.^4^

Circadian timing adds a systems-level constraint to these pathways. The mammalian circadian system organizes daily oscillations in transcription, metabolism, redox state, DNA repair capacity, and cellular stress responsiveness, defining time-of-day windows that differ in permissiveness for synthesis, repair, degradation, and recovery.^5,6,7^ Importantly, circadian control is not limited to transcriptional programs: post-translational regulation and controlled protein turnover are central to clock precision, entrainment, and robustness.^7^ In this context, a PAR-linked ubiquitin layer becomes particularly consequential. Because PAR synthesis draws directly on NAD⁺, and because NAD⁺ metabolism is tightly coupled to circadian physiology and behavioral state, PAR generation is positioned to be temporally structured rather than uniform across the day.^5,6,7^ If PAR availability fluctuates across time-of-day windows, then PAR-binding proteins, particularly PAR-responsive E3 ligases, become candidate effectors that translate circadian context and stress intensity into phase-dependent proteostatic outcomes.^7,8^

Neurodegenerative disorders provide a setting in which these intersections are likely to be functionally important. PAR signaling and PAR-binding effectors have been repeatedly implicated in neurodegenerative pathways—particularly in proteostasis- and stress-linked contexts—yet this biology is rarely framed through a circadian lens, even though circadian and sleep disruption are pervasive across neurodegenerative disease. In parallel, clock robustness depends heavily on post-translational regulation and precisely timed protein turnover, creating a direct mechanistic opening for a PAR-responsive ubiquitin layer to influence circadian state. The PAR signalosome provides a rapid upstream stress-activated platform; PAR-binding E3 ligases provide substrate-selective control over ubiquitin flux and protein fate; and circadian timing defines structured windows in which these ubiquitin decisions can be most consequential for clock dynamics. Together, these layers motivate a coupled framework in which PAR-dependent ubiquitination is positioned to modulate core circadian regulation, such that disruption or mis-timing of PAR–E3 coupling could bias neural systems toward circadian rhythm instability in disease-relevant contexts.

Here, our goal is to determine whether non-covalent PAR-binding E3 ubiquitin ligases constitute a brain-deployed effector layer capable of coupling PAR signaling to circadian timing and disease-relevant proteostasis. To do so, we integrate GTEx v10 brain expression^9^, GWAS Catalog gene-mapped associations, CIRCA phase annotations^10^, and Human Protein Atlas cell-type transcriptomics^11^ to test. This work is driven by four questions: (i) what is the regional expression architecture of the PAR-linked E3 repertoire across human brain tissues; (ii) does human genetics for neurodegenerative, cognitive, and sleep/circadian phenotypes converge preferentially on PAR-binding ligases; (iii) is the PAR-binding subset non-randomly structured across circadian phase relative to non-PAR ligases, consistent with time-of-day “readout” of transient PAR availability; and (iv) is PAR-linked ubiquitin capacity cell-type weighted in a manner that implies differential interpretation of PAR pulses across neural populations.

Together, these analyses establish a systems-level framework for a non-covalent PAR–E3 ligase–circadian axis in neural tissues and define prioritized candidates and testable predictions for downstream validation.

## Methods

### Definition of the E3 ubiquitin ligase genes

To establish a consistent and biologically meaningful baseline for all downstream analyses, we first defined a comprehensive universe of human E3 ubiquitin ligases. We curated a set of 614 E3 ligases based on the presence of canonical E3-associated domains and motifs (including RING, HECT, RBR, and SCF-associated components), drawing from curated domain annotations and prior literature defining ubiquitin ligase architecture.^3^ Gene identifiers were harmonized to HGNC-approved gene symbols, and ambiguous aliases or deprecated identifiers were resolved manually to ensure one-to-one mapping across datasets. This 614-gene set was treated as the core ubiquitin “decision layer” of the proteostasis network, reflecting enzymes that confer substrate selectivity and contextual control over ubiquitination. All subsequent analyses, including expression profiling, genetic enrichment, circadian phase mapping, and cell-type weighting, were performed within this defined universe to avoid biases arising from genome-wide background comparisons.

### Curation of PAR-binding E3 ligase sets

To operationalize PAR responsiveness within the E3 ligase layer, we defined a PAR-binding motif-positive E3 subset using the supplemental E3 ligase analysis reported by Kang et al.^3 From the reference set of 614 human E3 ubiquitin ligases, 327 unique ligases contained at least one of three PAR-binding motif classes: the newly identified CPxC/CNxC motif or either of two previously defined PAR-binding motifs (Table S14 of Kang et al.^3). This 327-gene set was used as the PAR-binding E3 ligase reference set for downstream expression, genetic, circadian, and cell-type analyses.

### GTEx v10 bulk transcriptome resources and brain tissue selection

Bulk tissue expression analyses used GTEx Analysis Release V10 RNA-seq expression quantifications (RNASeQC v2.4.2; dbGaP accession phs000424.v10.p2). First, the tissue-level median TPM matrix (GTEx_Analysis_v10_RNASeQCv2.4.2_gene_median_tpm.gct.gz) was used for tissue-wide summaries and for defining brain-deployed/background E3 universes.

Second, sample-level TPM values (GTEx_Analysis_v10_RNASeQCv2.4.2_gene_tpm.gct.gz) were used for region-resolved distribution plots; sample identifiers in this matrix were mapped to detailed tissue labels using the GTEx sample attribute table (GTEx_Analysis_v10_Annotations_SampleAttributesDS.txt). For brain-region-specific analyses, we restricted attention to the 13 GTEx brain regions (tissues labeled Brain_*), and for sample-level plots, we retained only samples annotated to those brain regions in the sample attribute table.^9

### Tissue-wide expression summaries and definition of brain detection

To summarize tissue-wide deployment of the E3 ligase layer (**Fig. 1A**), GTEx v10 tissue-level median TPM values were reshaped to long format and summarized per tissue across the fixed 614-ligase universe. Two tissue-level metrics were computed for each tissue: (i) the median-of-medians, defined as the median TPM across the 614 ligases within that tissue, and (ii) the detection rate, defined as the fraction of ligases with median TPM ≥ 1 within that tissue. Tissue labels were standardized by converting underscores to spaces, and tissues were ordered by median-of-medians to preserve identical ordering across summary panels **(Supplementary Table 2)**. For downstream genetic analyses, we defined a brain-deployed background set of ligases using GTEx v10 tissue-level median TPM across brain regions (13 tissues labeled Brain_*, as defined in GTEx v10 bulk transcriptome resources and brain tissue selection). A ligase was classified as brain-detected if its median TPM was ≥ 1 in at least one brain region, yielding a background brain-detected E3 set of 483 ligases **(Supplementary Table 1).** To support downstream analyses requiring a brain-enriched subset, the precomputed Top 483 brain-enriched ligase list was appended to the 614-ligase master table as a binary membership flag and distributed as part of Supplementary Table 1, which served as the authoritative reference for subsequent regional and cell-type analyses.

**Figure 1.**
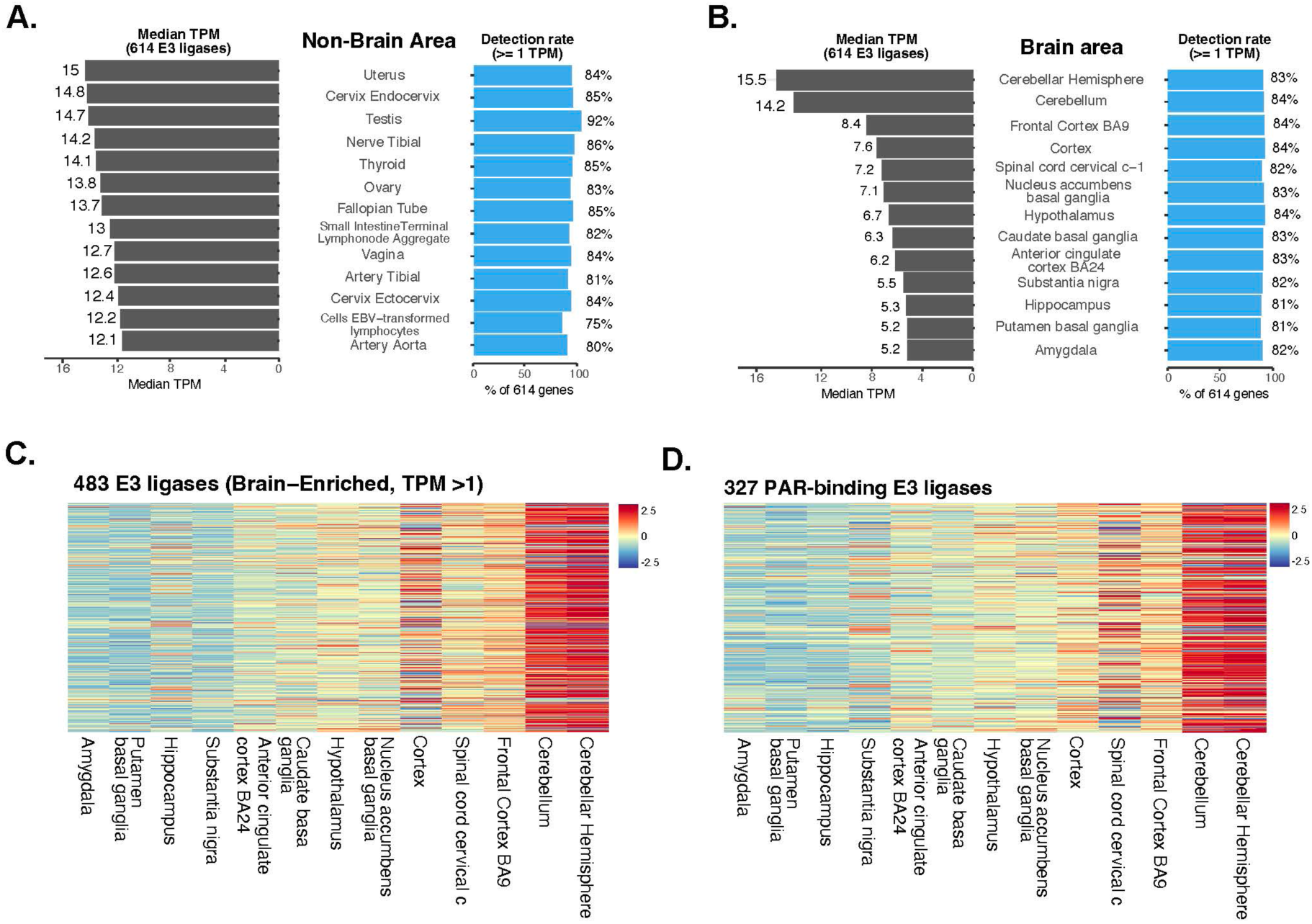
Brain-wide deployment and regional organization of PAR-binding E3 ubiquitin ligases. (A) Median expression (left; TPM) and detection rates (right; ≥1 TPM) for the 614-reference E3 ligase set across the top 13 non-brain GTEx tissues ranked by median E3 abundance. Detection rates remain consistently high across tissues despite organ-to-organ variation in median expression. (B) Median expression (left; TPM) and detection rates (right; ≥1 TPM) across the 13 GTEx brain regions. E3 ligase expression is broadly maintained across CNS compartments, with comparatively higher median abundance in cerebellar and cortical tissues. (C) Row-normalized heatmap (z-score) of the 483 brain-enriched E3 ligases (TPM >1) across GTEx brain regions, demonstrating regional enrichment structure within the brain E3 landscape. (D) Row-normalized heatmap (z-score) of the 327 PAR-binding E3 ligases retained after brain-enrichment filtering. The PAR-binding subset preserves the broader regional organization observed in the larger brain-enriched E3 repertoire. Supplementary Tables 1–3 provide underlying expression and detection values.

### Brain regional heatmap construction using the Top 483 brain-enriched ligase set

To visualize regional structure within the brain-enriched E3 layer (**Fig. 1B–C**), we used the precomputed Top 483 brain-enriched ligase set encoded in the master ligase table **(Supplementary Table 1).** For heatmap generation, GTEx v10 tissue-level median TPM values were subset to the Top 483 ligases and to the 13 GTEx brain-region columns (Brain_*). A genes × brain-regions matrix was assembled from median TPM values, with missing values filled as 0.

To stabilize low-abundance values prior to normalization, TPM was log-transformed as (*TPM* + 0.1), where 0.1 prevents undefined values at TPM = 0. Heatmaps were plotted using row-wise z-score normalization (each gene standardized across brain regions) to emphasize relative regional enrichment patterns independent of absolute expression magnitude. Brain-region columns were ordered deterministically by the median TPM across the Top 483 genes (lowest to highest), and clustering was disabled (cluster_rows = FALSE; cluster_cols = FALSE) to preserve a fixed region order across panels and facilitate direct visual comparison.

### Representative ligase selection strategy and violin plotting

To document the basis for representative selection (**Fig. 2A**), we positioned all Top 483 ligases on a two-parameter map computed from GTEx brain-region median TPM values: (i) the median TPM across the 13 brain regions and (ii) the standard deviation of the 13 region-specific medians. Representatives were selected to span this space so that the violin panels reflect distinct abundance/variability regimes rather than a single expression archetype **(Supplementary Table 3B)**.

**Figure 2.**
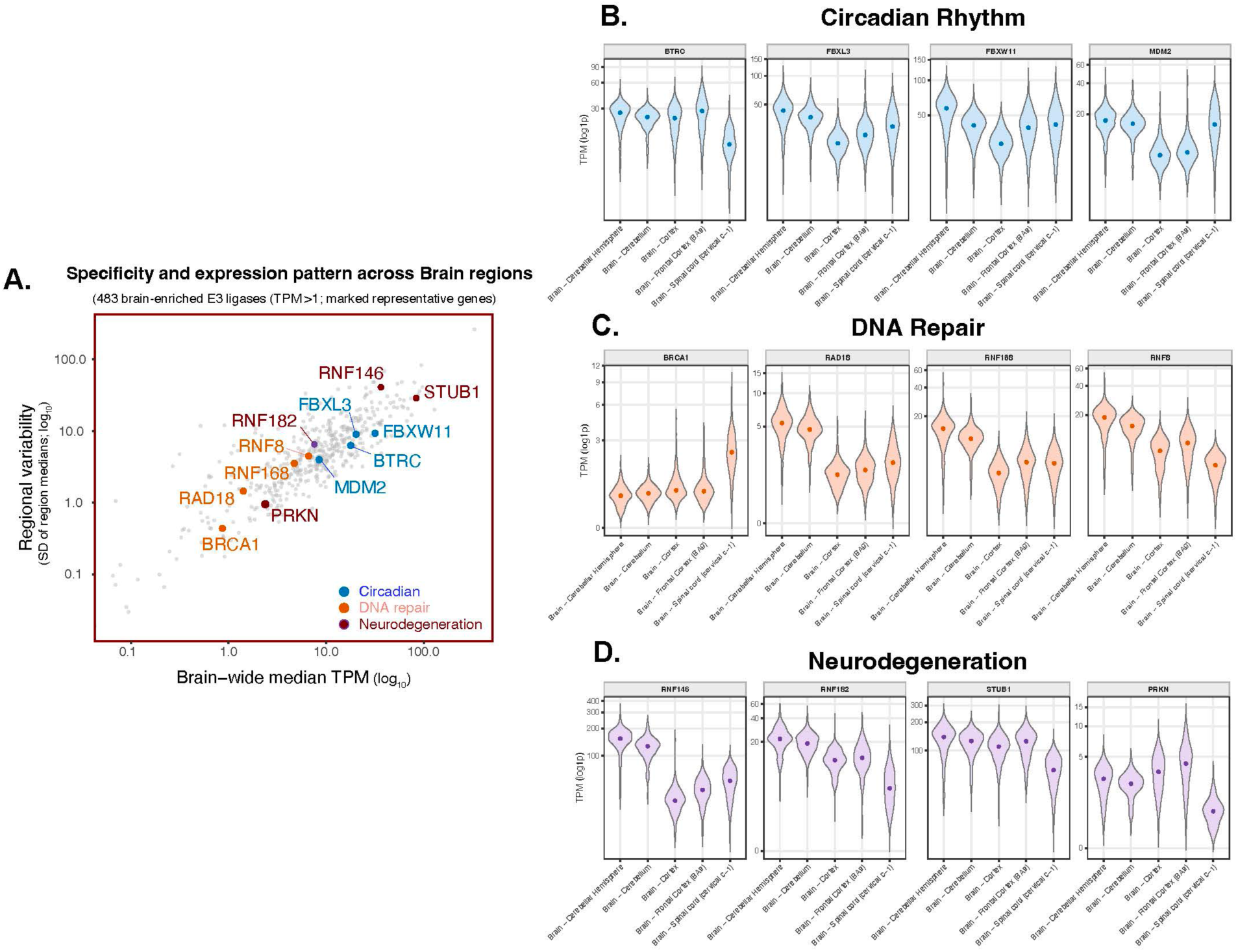
Representative expression archetypes within the brain-enriched E3 ligase repertoire. (A) Selection map showing representative ligases positioned by brain-wide median TPM (x-axis; log10) and regional variability across GTEx brain regions (y-axis; SD of log10-transformed regional median TPM values). Representative ligases were selected to span distinct abundance and variability regimes within the broader set of 483 brain-enriched E3 ligases. (B) Representative circadian-linked ligases (BTRC, FBXL3, FBXW11, MDM2) display broadly distributed expression across brain regions with moderate regional variation. (C) Representative DNA repair ligases (BRCA1, RAD18, RNF168, RNF8) exhibit comparatively low-but-constitutive expression across brain regions. (D) Representative neuronal stress/proteostasis and neurodegeneration-linked ligases (RNF146, RNF182, STUB1, PRKN) show higher-amplitude and/or more regionally structured expression patterns. Violin plots represent GTEx v10 sample-level TPM distributions across representative brain regions (log1p TPM). Full 13-region distributions are provided in Supplementary Fig. 1A–C.

To generate interpretable examples of expression behavior within the brain-enriched E3 layer (**Fig. 2B–D**), we selected twelve representative ligases spanning circadian-linked regulation, DNA repair, and neuronal stress/proteostasis pathways. Representatives were chosen to cover a range of brain-wide abundance and regional variability within the Top 483 set rather than to optimize any single metric; PRKN was retained as an explicit example of cell-type–restricted expression that can be underestimated in bulk tissue. Violin plots were constructed from GTEx v10 sample-level TPM values. Samples were assigned to brain regions by mapping sample identifiers in the TPM matrix to GTEx detailed tissue labels using the sample attribute table, then restricting to the 13 GTEx brain regions. For each selected ligase, the distribution of sample-level TPM across brain regions was visualized as a violin plot after applying a log-stabilizing transform: Violin y-axis transforms: *log log* (1 + *TPM*) **(Supplementary Table 3A).** For readability, the main **Fig. 2** panels display a representative subset of GTEx brain regions, whereas full 13-region distributions are provided in **Supplementary Fig. 1A–C**.

### GWAS Catalog acquisition, gene mapping, and PAR-binding enrichment tests

To test whether PAR-binding E3 ligases are preferentially implicated in neurological and sleep/circadian phenotypes, we analyzed GWAS Catalog associations using a gene-mapped workflow and generated both (i) umbrella-level overlap counts for visualization and (ii) a standardized gene-by-umbrella matrix for majority testing. GWAS associations were obtained from the GWAS Catalog bulk associations table (gwas_catalog_v1.0.2-associations_e115_r2025-09-29.tsv) downloaded from the GWAS Catalog FTP “latest releases” directory.^12^ Trait names were taken from the “DISEASE/TRAIT” field. Gene mapping was defined directly from GWAS Catalog annotations by extracting the union of REPORTED GENE(S) and MAPPED_GENE fields for each association, splitting multi-gene strings on standard delimiters (commas/semicolons), trimming whitespace, removing non-informative placeholders (e.g., “NR”), and harmonizing symbol aliases where needed (e.g., PARK2→PRKN) to match the ligase annotation tables. Associations were restricted to ACC-relevant phenotypes using keyword-based rules capturing sleep/circadian and neurodegeneration/cognition–related terms, and filtered traits were assigned to ten umbrella phenotypes using deterministic string-matching rules: Cognitive Performance, AD, Insomnia, Circadian Rhythm, Stroke, PD, Age-Related Cognitive Decline, ALS, FTD, and HD **(Supplementary Table 4A)**. For each umbrella phenotype, we summarized the GWAS signal as the number of unique associated E3 ligases after intersecting GWAS-mapped genes with the annotated ligase universes in **Supplementary Table 1**: the brain-detected background (n = 483; defined using GTEx v10 median TPM across the 13 GTEx brain regions) and the PAR-binding motif-positive set (n = 327). These overlap counts were visualized as two-bar summaries per umbrella phenotype

To enable reproducible majority testing, we additionally constructed a wide gene-by-umbrella membership matrix (ligases_traits.csv) from the same filtered, umbrella-assigned GWAS associations by collapsing to unique gene–umbrella pairs and pivoting to boolean umbrella columns (TRUE if a gene had ≥1 association assigned to that umbrella; FALSE otherwise) **(Supplementary Table 4B).** Majority testing was performed within the brain-detected background: for each umbrella phenotype, n was the number of brain-detected trait-linked E3 ligases, and k was the number of those ligases annotated as PAR-binding motif-positive. We reported Wilson 95% confidence intervals for k/n and performed one-sided exact binomial tests against a 50% reference (H0: p = 0.5; H1: p > 0.5). Reported p-values are nominal and are interpreted as descriptive evidence of directionality across phenotypes rather than as strict multiple-testing–corrected significance.

### Circadian phase annotation and processing

To restrict analyses to rhythms consistent with circadian periodicity, entries were filtered to a near-24 h band:

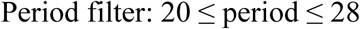

Because CIRCA^10^ may report multiple phase calls for the same gene across different studies and tissues, we prevented over-weighting of genes with many entries by retaining a single representative phase call per ligase under a consistent one-call-per-gene rule for class-level comparisons. Phase organization was summarized by binning *p*ℎ*ase*_24_ into integer hour bins (0– 23) and comparing the resulting per-bin ligase counts between PAR-binding and non-PAR ligase classes using identical binning and plotting parameters.

To interpret timing relative to the core-clock reference state rather than absolute clock time, we computed phase offsets relative to BMAL1 and CRY1. For each ligase and reference gene, offsets were computed as a wrapped difference:

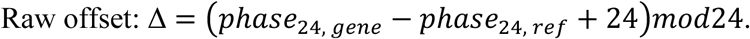

Offsets were then mapped to a symmetric interval to represent leads and lags around the oscillator:

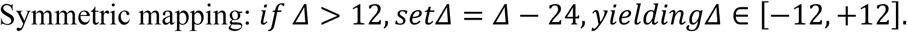

Offset distributions for PAR-binding versus non-PAR ligases were compared using consistent binning and visualization settings. Circular phase plots were used as qualitative visualizations of phase placement across the 24-hour cycle, providing a bin-free view consistent with the phase-density and offset summaries.

### Cell-type specific expression analysis using the Human Protein Atlas

To evaluate whether circadian-linked E3 ubiquitin ligases, and specifically the PAR-binding circadian subset, are differentially weighted across human brain cell types, we performed a program-level expression enrichment analysis using the Human Protein Atlas (HPA)^11^ single-cell type transcriptomics resource. HPA reports normalized expression as nCPM aggregated across curated cell types; these values are library-size normalized within the HPA framework and therefore support within–cell type comparisons of gene-level expression. All analyses were restricted to the Top 483 brain-enriched E3 ligases to maintain a fixed baseline “E3 program” across cell types and to ensure that enrichment scores reflect allocation within the brain-deployed ubiquitin ligase layer rather than genome-wide differences **(Supplementary Table 6A)**.

Let C denote a given brain cell type, and let GE3 denote the Top 483 E3 ligase universe. For each cell type C, we defined the total E3 expression program as the summed expression mass across all E3 ligases in the baseline set:

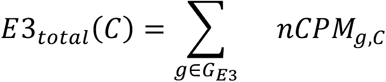

For a target gene subset S ⊆ GE3, (e.g., circadian-linked E3s or PAR-binding circadian E3s), we computed its observed expression mass and its observed share of the total E3 program:

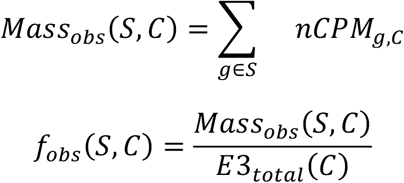

To normalize for gene-set size (i.e., to prevent larger sets from appearing enriched simply because they contain more genes), we defined a gene-count null expectation for the subset share within the E3 universe:

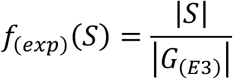

We then computed an observed-versus-expected enrichment ratio and plotted it on a log scale. Specifically, for each cell type C and subset S, the enrichment ratio was:

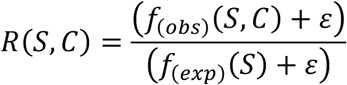

where ε is a small pseudocount included solely to avoid undefined values when *f_obs_*(*S*, *C*) approaches zero. The final reported score was the log-transformed enrichment ratio:

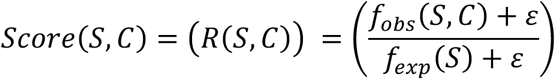

This formulation yields a directly interpretable program-level enrichment metric with a natural null. A score of 0 indicates that the subset contributes exactly its gene-count–expected fraction of the E3 program in that cell type; positive values indicate enrichment beyond expectation (e.g., +1 corresponds to ∼2-fold enrichment), and negative values indicate depletion (e.g., −1 corresponds to ∼2-fold depletion). Because both observed and expected quantities are expressed as fractions of the same baseline E3 program, this metric is robust to differences in overall transcriptional output across cell types and reflects how strongly a given cell type allocates E3 expression toward the subset of interest.

We applied this identical computation to two subsets. First, we defined S = Gcirc, the circadian-linked E3 ligases, to quantify circadian E3 program enrichment per cell type. Second, we defined S = GPAR-circ ⊂ Gcirc, the PAR-binding circadian-linked E3 ligases, to quantify enrichment of the PAR-responsive circadian E3 program. In both cases, enrichment was assessed within the fixed Top 483 E3 universe and anchored to E3total(C), enabling direct comparison of circadian and PAR-responsive circadian program weighting across neuronal and glial populations (**Supplementary Table 6B**, complementary within-circadian PAR-share summaries in **Supplementary Table 6C**).

### Statistical analysis and visualization

All analyses were performed in R using standard tidyverse workflows. Heatmaps, violin plots, bar plots, phase-density plots, and circular visualizations were generated using ggplot2 and associated packages. Statistical tests are specified in the relevant sections above. No data points were excluded post hoc, and all analyses were performed within pre-defined gene universes.

## Results

### Regional organization of brain-enriched and PAR-binding E3 ubiquitin ligases

Before mapping PAR-binding ligases across brain tissues, we first established the baseline architecture of the E3 ubiquitin ligase layer itself. E3 ligases are the substrate-selective “decision makers” of the ubiquitin–proteasome system, and in the brain, even modest shifts in E3 deployment can reshape long-term proteostasis, repair complex turnover, synaptic stability, mitochondrial quality control, and inflammatory signaling. Accordingly, we asked where the broader E3 repertoire is present in the CNS, whether that deployment is regionally structured, and what fraction of the brain-enriched atlas is positioned to interface with PAR through non-covalent binding.

Using GTEx v10 transcriptomes, we summarized tissue-wide expression and detection for our reference set of 614 E3 ligases; 606/614 mapped to GTEx v10 gene symbols (8 absent or unmapped) and were analyzed here. Across tissues, most ligases are detectable despite organ-to-organ differences in median expression, indicating that this E3 layer is broadly deployed rather than restricted to a narrow subset of compartments. To provide a broader tissue context, we first summarized expression across the top 13 non-brain tissues ranked by median E3 abundance (**Fig. 1A, Supplementary Table 2).** Across these tissues, median E3 expression and detection rates remain consistently high, indicating that the E3 ligase layer represents a widely deployed regulatory system rather than a compartment-restricted program.

We next focused specifically on CNS deployment by examining expression across the 13 GTEx brain regions (**Fig. 1B, Supplementary Table 2).** Within the brain, E3 ligase signals are broadly maintained across regions rather than being dominated by a single anatomical compartment. Cerebellar and cortical tissues exhibit among the highest median expression levels, whereas several basal ganglia-associated regions show comparatively lower median abundance; however, detection rates remain consistently high across all sampled brain regions. Together, these results indicate that the E3 ubiquitin ligase layer is broadly deployed throughout the CNS while retaining modest regional structure.

To resolve regional structure within the brain-enriched E3 layer, we next examined the top brain-enriched E3 ligase set (n = 483) across GTEx brain regions (**Fig. 1C, Supplementary Table 1).** The brain-enriched atlas shows clear structure, with broad relative enrichment across cerebellar and cortical compartments for large subsets of ligases and comparatively lower relative enrichment across several basal ganglia and midbrain-associated regions. Importantly, because the heatmap is row-normalized (z-scores), these patterns reflect regional enrichment structure rather than absolute expression magnitude. Lastly, we connected this expression architecture to PAR biology by restricting to ligases containing at least one of the defined PAR-binding motifs. Of 327 PAR-binding E3 ligases in our reference list, 321 were retained in the GTEx brain atlas after symbol matching and brain-enrichment filtering, yielding a PAR-binding brain-enriched interactome (n = 321) (**Fig. 1D).** This PAR-linked atlas preserves the same broad regional logic observed in the larger brain-enriched set, emphasizing that PAR-binding is layered onto an E3 program that is both widely deployed and regionally structured across the brain.

### Representative regional expression archetypes across circadian, DNA repair, and neurodegeneration-linked E3 ligases

To highlight the range of expression behaviors embedded within the brain-enriched E3 ligase layer, we first positioned twelve representative ligases within the broader brain-enriched E3 landscape using abundance and regional-variability metrics (**Fig. 2A**). These ligases were selected to span three mechanistic categories: circadian-linked ubiquitin regulation (BTRC, FBXL3, FBXW11, MDM2), DNA repair ubiquitin signaling (BRCA1, RAD18, RNF168, RNF8), and neuronal stress/proteostasis and neurodegeneration-relevant pathways (RNF146, STUB1, RNF182, PRKN) (**Fig. 2B–D**). For readability, Figure 2 displays a representative subset of brain regions, while the full 13-region GTEx distributions for all plotted ligases are provided in **Supplementary Fig. 1A–C**.

To make the representative selection transparent, we positioned these twelve ligases within the broader Top-483 brain-enriched E3 landscape using a two-parameter selection map defined by brain-wide median TPM (x-axis; log10) and regional variability across GTEx brain regions (y-axis; SD of log10-transformed regional median TPM values) (**Fig. 2A, Supplementary Table 3B)**. The representatives occupy distinct regions of this space rather than clustering into a single regime: circadian-linked ligases largely populate a moderate-to-high abundance band; DNA repair ligases cluster toward lower abundance and lower variability; and neuro/proteostasis ligases extend into higher abundance and/or higher regional heterogeneity. This distribution emphasizes that the representative panels were selected to capture multiple expression archetypes embedded within the broader brain-enriched E3 repertoire rather than a single expression profile.

Circadian-linked ligases show broad and generally uniform expression across brain regions, with moderate-to-high distributions across cortex, hippocampus, and cerebellar compartments (**Fig. 2B, Supplementary Table 3A)**. Across these examples, cerebellar and cortical regions frequently sit toward the higher end of the regional medians, whereas multiple basal ganglia– associated regions trend lower. This pattern should be interpreted strictly as a bulk regional transcript signature, not a direct readout of circadian “activity”, but it is consistent with widespread neural deployment of clock-adjacent ubiquitin machinery. Mechanistically, these representatives map onto established ubiquitin control points in clock protein turnover, including SCF(FBXL3) regulation of CRY stability,^13,14^ β-TrCP-dependent control of PER2 turnover,^15^ and MDM2-mediated regulation of PER2 stability and period control.^16^

DNA repair ligases (BRCA1, RAD18, RNF168, RNF8) follow a distinct “low-but-constitutive” bulk expression archetype across brain regions (**Fig. 2C).** Absolute levels are lower than many circadian-linked and proteostasis examples, but detectability is maintained across the sampled compartments, including cortex and hippocampus, with comparatively modest region-to-region shifts. This is consistent with the expectation that core ubiquitin-dependent DNA damage signaling is maintained as a baseline cellular program rather than being confined to a narrow neuroanatomical compartment. The selected representatives anchor well-characterized nodes in DSB-linked ubiquitin signaling, including RNF8-dependent lesion ubiquitylation and recruitment cascades,^17^ RNF168 amplification/ubiquitin decoding functions,^18^ and RAD18 integration of DNA damage signaling with homologous recombination programs.^19^

Neuronal stress/proteostasis and neurodegeneration-relevant ligases show higher amplitude and/or clearer regional structure (**Fig. 2D**). RNF146 and STUB1 are broadly and robustly expressed, with especially high distributions in cerebellar and cortical compartments, consistent with their established roles in PAR-linked ubiquitin signaling (RNF146/Iduna)^4^ and chaperone-coupled proteostasis control (CHIP/STUB1).^20^ RNF182 shows preferential enrichment toward cortical and hippocampal regions, aligning with prior reports describing RNF182 as a brain-enriched E3 ligase up-regulated in Alzheimer’s disease brain tissue.^21^ By contrast, PRKN displays comparatively low bulk expression across GTEx brain regions, including substantia nigra (**Fig. 2D).** Given that bulk RNA-seq reflects tissue mixtures and can dilute signals from region- and cell-type-restricted programs, we treat PRKN here as an explicit control for this interpretive limitation rather than as an absence-of-expression claim; notably, Parkin protein expression has been reported across multiple CNS regions in immunohistochemical studies.^22^

Together, Figure 2 illustrates three recurring expression archetypes within the brain-enriched E3 repertoire: (i) broadly expressed clock-adjacent ligases with moderate-to-high abundance, (ii) repair-associated ligases with low-but-constitutive bulk expression, and (iii) stress/proteostasis ligases with higher amplitude and/or clearer regional structure. These representative patterns provide interpretable anchors for the genetic and circadian timing analyses that follow.

### PAR-binding E3 ligases are enriched among GWAS-linked neurological and circadian traits

Having established that PAR-linked ligases are broadly expressed across neural tissues with enrichment in cortical and cerebellar compartments, we next asked whether human genetics converges on this same subset. Specifically, we tested whether E3 ligases mapped to genome-wide association study (GWAS) loci for Alzheimer’s disease (AD), cognition, circadian traits, and related neurological phenotypes are selectively enriched for PAR-binding activity. This analysis is motivated by a mechanistic premise: PAR is not only a covalent modification, but also a high-avidity scaffold that signals through non-covalent PAR “reader” interactions; therefore, E3 ligases that can bind PAR are positioned to convert transient PAR pulses into selective ubiquitination programs, providing a plausible route by which PAR signaling can interface with circadian timing and neurodegenerative vulnerability.

We compared three gene sets that separate “brain deployment” from “PAR responsiveness”. First, we defined a brain-detected background of ligases (n = 483; GTEx v10 median TPM ≥ 1 in ≥ 1 brain region).^9^ Second, we used a PAR-binding motif-positive set (n = 327) defined from the E3 ligase motif analysis of Kang et al.^3^ Using these parallel universes allows us to test whether genetic signals track specifically with PAR-binding capacity, rather than simply reflecting broad E3 expression in the brain.

Trait signals were aggregated under ten ACC-relevant umbrella phenotypes using large GWAS studies: cognitive performance²³; AD²⁴,²⁵; insomnia²⁶; chronotype/circadian rhythm²⁷; stroke²⁸; Parkinson’s disease²⁹; general cognitive function³⁰; ALS³¹; FTD³²; and HD modifiers³³ (Supplementary Table 4A). For each umbrella, we intersected GWAS-mapped E3 ligases with the three gene sets above and summarized counts (**Fig. 3A**). In the highest-signal phenotypes (cognitive performance, AD, insomnia, circadian rhythm, stroke, and PD), the PAR-binding subset closely tracks or exceeds the broader 483-ligase background. Practically, this indicates that, where GWAS implicates E3 ligases in these phenotypes, a substantial fraction of those ligases are PAR-binding, as supported by curated evidence.

**Figure 3.**
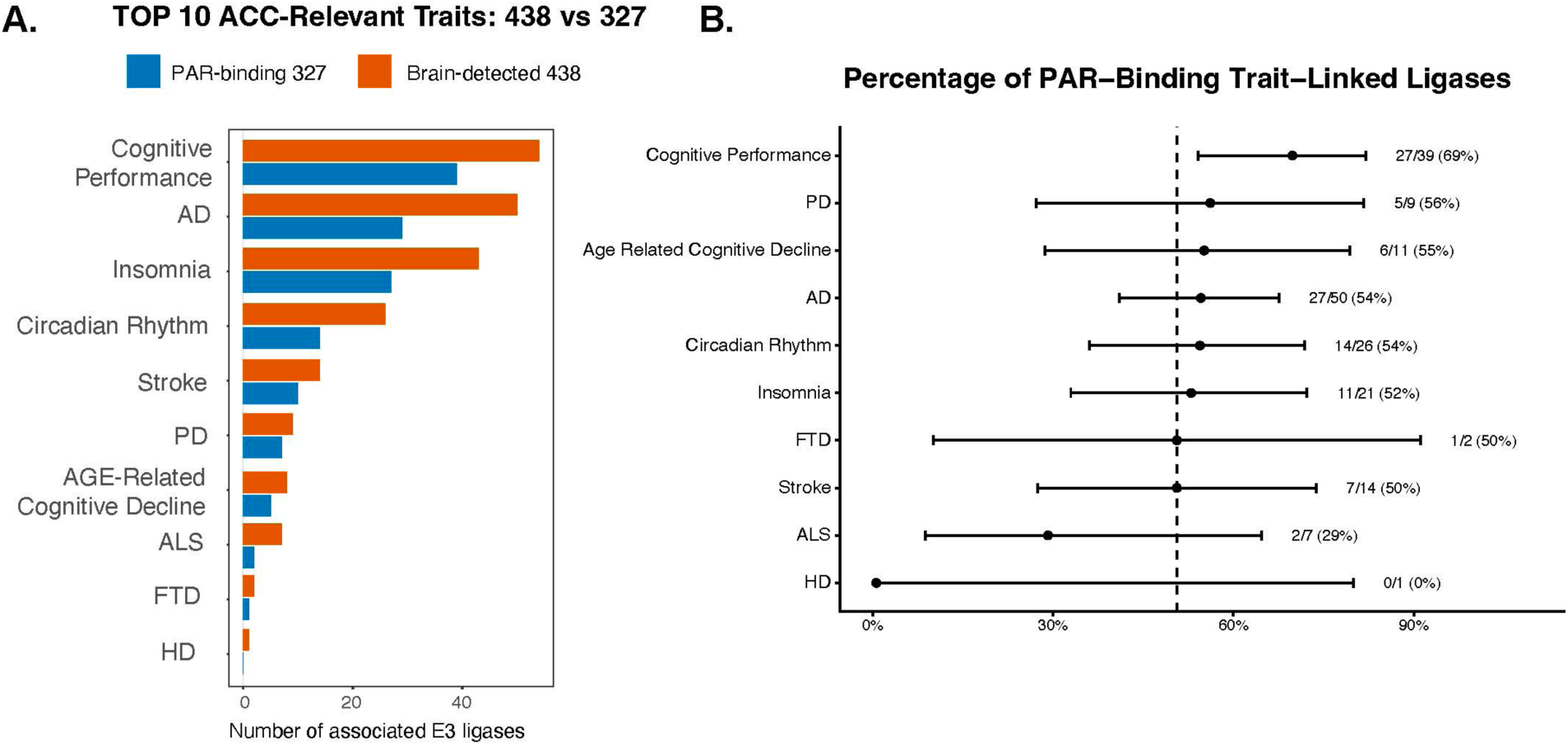
Genetic convergence of PAR-binding E3 ligases across ACC-relevant phenotypes. (A) Number of E3 ligases associated with the top ACC-relevant GWAS umbrella phenotypes, comparing the broader brain-detected E3 set (n = 438) with the PAR-binding subset (n = 327). Traits include cognition-, neurodegeneration-, and circadian/sleep-related categories. (B) Percentage of trait-linked E3 ligases that are PAR-binding across the same phenotypes. Values indicate the proportion of GWAS-associated ligases within each trait category that possess predicted or experimentally supported PAR-binding capacity. Dashed line indicates the background proportion of PAR-binding ligases within the analyzed brain-detected E3 repertoire.

To quantify whether PAR-binding ligases constitute a majority of trait-linked E3 hits, we computed for each umbrella the fraction of trait-linked ligases that are PAR-binding and plotted Wilson 95% confidence intervals, using one-sided exact binomial tests against a 50% reference (**Fig. 3B, Supplementary Table 4B).** Cognitive performance shows the strongest shift in this panel (69% PAR-binding; p = 0.0119), while AD, PD, insomnia, and circadian rhythm trend above the majority threshold with more modest effect sizes (**Fig. 3B).** Together, these results support a systems-level convergence: across cognition-, sleep/circadian-, and neurodegeneration-associated phenotypes, GWAS-linked E3 ligases are disproportionately PAR-responsive, consistent with a model in which PAR-dependent ubiquitination represents a central interface connecting stress signaling, proteostasis, and circadian homeostasis.

### Circadian phase organization of PAR-binding E3 ubiquitin ligases

To test whether PAR-binding capacity is associated with circadian timing structure, we leveraged circadian phase annotations from CircaDB/CIRCA, which compiles mammalian time-course expression datasets and reports rhythm statistics, including phase, period, and significance metrics derived from rhythm-detection algorithms such as JTK_Cycle and related approaches.^10^ Each ligase is assigned a peak phase (in hours) along with periodicity measures, and we stratified ligases by curated PAR-binding status (PAR-binding vs non-PAR) to ask whether PAR-binding ligases occupy structured circadian windows relative to the broader E3 repertoire **(Supplementary Table 5).**

Because CIRCA integrates phase calls across multiple experiments (and therefore the same gene can have slightly different reported phases depending on tissue context, entrainment conventions, and rhythm-calling thresholds), we standardized phase representation prior to class comparisons. We wrapped the phase into a 24-hour interval (phase mod 24) and restricted it to a near-24 h periodicity band (period 20–28 h) to focus on rhythms plausibly aligned with circadian time. To avoid over-weighting genes with multiple entries, we retained a single representative phase call per ligase, applying a consistent one-call-per-gene selection rule, yielding one phase per ligase for downstream comparisons **(Supplementary Table 5).**

We first summarized the timing structure using a phase-density view. Peak phases were binned into integer-hour bins (0–23), and we counted the number of ligases peaking in each bin separately for the PAR-binding and non-PAR classes. In this phase map, PAR-binding ligases show clear non-uniformity with discrete windows of enrichment and depletion rather than an even distribution across the cycle (**Fig. 4A).** The non-PAR class displays a lower-amplitude and differently shaped density profile, indicating that the PAR-binding timing structure is not simply a scaled version of the non-PAR background.

**Figure 4.**
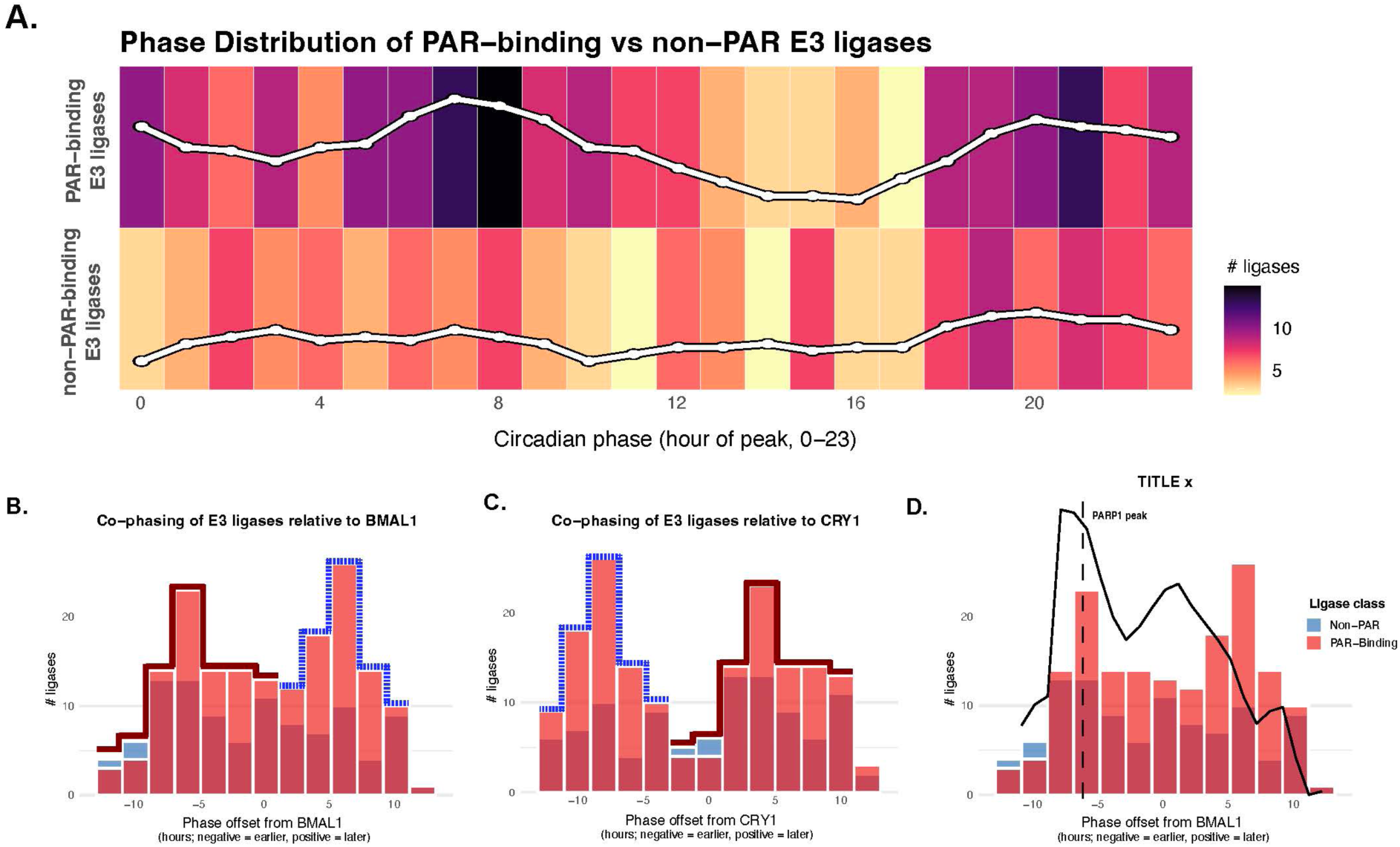
Circadian phase organization of PAR-binding E3 ubiquitin ligases. (A) Circadian phase distributions of PAR-binding and non–PAR-binding E3 ligases across the 24-hour circadian cycle using CIRCA phase annotations. Heatmap intensity and overlaid curves indicate the number of ligases peaking at each circadian phase. (B) Distribution of circadian phase offsets for E3 ligases relative to BMAL1 peak phase. Negative values indicate earlier peak timing relative to BMAL1; positive values indicate later timing. (C) Distribution of circadian phase offsets for E3 ligases relative to CRY1 peak phase. (D) Relative phase organization of PAR-binding and non–PAR-binding ligases around the reported PARP1 peak phase window, highlighting enrichment of PAR-binding ligases within circadian transition regions.

Because absolute circadian time can shift across experiments depending on entrainment and reference conventions, we next expressed timing relative to canonical core-clock anchors rather than relying solely on absolute phase. We computed each ligase’s phase offset relative to BMAL1 and CRY1, and compared the resulting offset distributions between PAR-binding and non-PAR ligases (**Fig. 4B–C).** Offsets were defined as a wrapped difference mapped onto a symmetric interval (e.g., −12 to +12 hours), so negative values indicate earlier-than-reference peak timing and positive values indicate later-than-reference timing.

In both reference frames, PAR-binding ligases show a clear, structured, bimodal co-phasing pattern rather than a uniform offset distribution (**Fig. 4B–C).** Relative to BMAL1, PAR-binding ligases exhibit one prominent lobe centered around approximately −6 hours, with a second emphasized lobe on the positive side (approximately +4 to +8 hours; highlighted by the dotted outlines in **Fig. 4B**). When the reference is shifted to CRY1, this same secondary concentration appears on the negative side (approximately −10 to −6 hours, again highlighted by the dotted outlines in **Fig. 4C**), consistent with the expected rotation of offsets when changing the anchor gene. In contrast, the non-PAR class remains comparatively lower-amplitude and less sharply structured across these offset windows (**Fig. 4B–C).** Together, these offset patterns indicate that PAR-binding ligases occupy preferred timing relationships around the oscillator rather than being randomly distributed across phase offsets.

Finally, to relate the E3-ligase phase organization to a PAR-production–relevant signal, we overlaid a PARP1 circadian expression waveform onto the BMAL1-centered phase-offset histogram (**Fig. 4D).** PARP1 time-course values (from CIRCA) were shifted to the same BMAL1-referenced coordinate system (BMAL1 = 0) and wrapped into a single 24-h cycle; the fitted PARP1 curve was then scaled to the histogram height for visualization (i.e., the y-axes are not directly comparable). In this alignment, the PARP1 peak falls within an offset window that also contains a noticeable concentration of PAR-binding ligases, and broader features of the PARP1 waveform span regions where PAR-binding counts are elevated relative to non-PAR ligases. We emphasize that this comparison is qualitative and correlational and is further limited by differences in source datasets (e.g., tissue context and measurement platforms), but it provides a visually intuitive way to contextualize the phase-density and offset results (**Fig. 4A–C**) alongside a representative PARP-linked rhythm without making strong causal claims.

### Cell-type organization of circadian-linked and PAR-binding E3 ligases

To move from regional brain structure (GTEx) and phase organization (CIRCA) into a cell-resolved context, we next asked whether the circadian-linked E3 program, and specifically its PAR-binding fraction, is differentially weighted across brain cell types. For this analysis, we used the Human Protein Atlas (HPA) single cell type transcriptomics resource, which provides normalized expression (nCPM) summarized across curated cell types from integrated scRNA-seq/snRNA-seq datasets.^11^ We restricted to HPA brain-relevant neuronal and glial classes and used the same baseline brain-enriched E3 universe (Top 483), enabling a direct “program share” view rather than single-gene anecdotes.

As a baseline control, we first plotted the mean nCPM across genes for each brain cell type (**Fig. 5A**). This panel is not intended to support a mechanistic claim; it simply establishes the expression scale and confirms that the subsequent enrichment metrics are not trivially driven by one cell type having uniformly higher transcript abundance across all genes. In other words, Figure 5A is the “burden context” that anchors the interpretation of the enrichment analyses.

**Figure 5.**
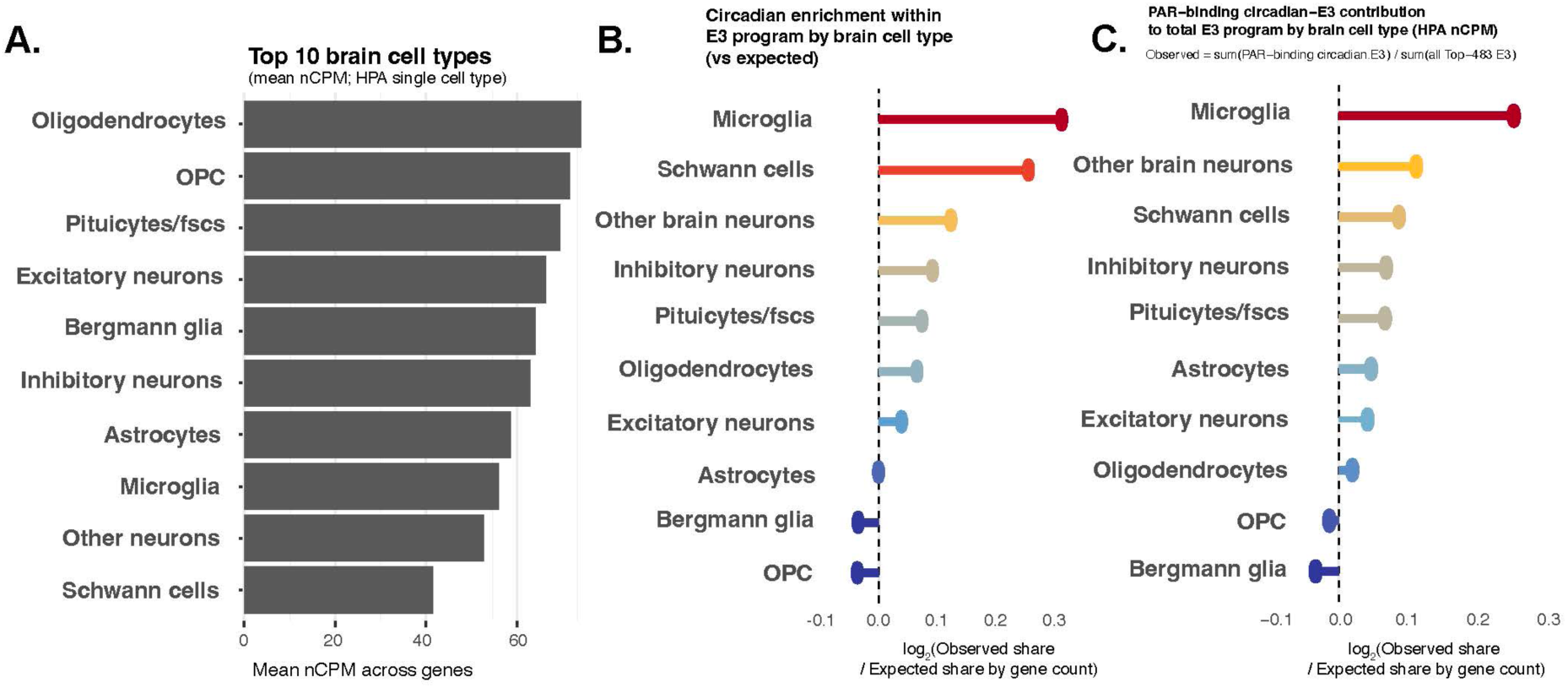
Cell-type weighting of circadian and PAR-binding circadian E3 programs in the brain. (A) Mean normalized expression (nCPM) of PAR-binding circadian E3 ligases across major Human Protein Atlas brain cell types. (B) Enrichment of circadian-linked E3 ligases within each brain cell-type E3 program relative to expectation by gene count. Values are shown as log2(observed share / expected share). (C) Relative contribution of PAR-binding circadian E3 ligases to the total E3 program within each brain cell type. Microglia show the strongest enrichment across both enrichment frameworks, supporting preferential weighting of PAR-responsive circadian ubiquitin regulation within microglial programs.

We then quantified whether circadian-linked E3 ligases account for more of the E3 program than expected from gene count alone. Specifically, we computed the observed circadian-E3 share within the baseline E3 set per cell type and normalized this by the expected share given the fraction of circadian-linked genes in the baseline set (**Fig. 5B).** Values above zero therefore indicate that circadian-linked E3 genes contribute more expression mass than expected purely by their membership fraction, whereas values below zero indicate relative depletion. In this analysis, circadian-linked E3 contribution is not uniform across brain cell programs: several cell types show positive deviation from expectation, while others fall at or below the null line (**Fig. 5B**). This supports the idea that circadian-linked ubiquitin control is not merely a constant fraction of the E3 layer but is modestly cell-type weighted in the brain. Lastly, we asked whether the PAR-binding fraction of the circadian-linked E3 layer exhibits additional cell-type structure beyond that captured by circadian weighting alone **(Supplementary Table 6C).** To capture this, we computed the observed contribution of PAR-binding circadian-E3 ligases to the total E3 program per cell type and again normalized by the expected share given gene count (**Fig. 5C, Supplementary Table 6B)**. This isolates a more specific question: not just “where are circadian E3s enriched,” but “where does the PAR-binding circadian subset disproportionately contribute to the overall E3 landscape.” In this view, several brain cell programs show positive deviation, indicating preferential weighting of PAR-binding circadian E3s, whereas others show relative depletion (**Fig. 5C).**

A key implication of these cell-type analyses is that microglia emerge as the dominant enrichment compartment for the circadian-linked E3 layer in this dataset. In the circadian-linked E3 share analysis (**Fig. 5B**), microglia rank highest above the gene-count expectation, indicating that circadian-linked E3 ligases contribute a disproportionately large fraction of the baseline E3 program in microglia relative to what would be predicted from set membership alone. This same pattern persists and sharpens when the analysis is restricted to the PAR-binding circadian subset normalized to the total E3 program (**Fig. 5C**), where microglia again show the strongest positive deviation. In other words, the two-step decomposition (circadian E3 weighting within the E3 program, followed by PAR-binding circadian-E3 weighting within total E3) points to microglia as the most consistently enriched brain cell program for a PAR-responsive circadian ubiquitin layer in the HPA nCPM resource.

This microglia-first signal has clear real-world relevance because microglia are the primary immune effector cells of the CNS and sit at the interface between circadian physiology and neurodegenerative vulnerability. Microglial inflammatory tone, phagocytic capacity, and synapse-associated clearance programs exhibit time-of-day dependence, and circadian disruption is repeatedly linked to exaggerated neuroinflammatory states and impaired clearance in disease-relevant contexts. In this framework, enrichment of circadian-linked E3 expression in microglia, and additional enrichment of the PAR-binding circadian subset, supports a mechanistic model in which PAR signaling can intersect microglial circadian state to bias ubiquitin-dependent regulation of inflammatory signaling, stress response pathways, and proteostatic turnover. While these data are correlational and do not measure ubiquitin activity directly, they identify microglia as the most strongly weighted compartment for the proposed PAR–E3–circadian axis and therefore a high-priority cellular context for downstream functional validation and translational timing considerations.

## Discussion

In this study, we propose and support a systems-level framework in which non-covalent PAR-binding E3 ubiquitin ligases constitute a temporally organized effector layer that couples PAR signaling to circadian timing and neural proteostasis. By integrating bulk tissue expression, human genetics, circadian phase annotations, and cell-type transcriptomic weighting, our analyses converge on a coherent picture: PAR-responsive ubiquitin control is not randomly distributed across tissues, genes, or time, but is structured across brain regions, circadian phase windows, and specific neural cell programs. This organization positions PAR-binding E3 ligases as plausible intermediates through which transient stress-activated PAR signals can be interpreted in a phase-dependent and cell-context–specific manner.

A key insight from the brain expression atlas is that the PAR-linked E3 repertoire is broadly deployed across the CNS while remaining regionally structured (**Fig. 1B–D).** The preservation of this regional structure upon restriction to PAR-binding ligases indicates that PAR responsiveness is layered onto an existing brain-deployed ubiquitin architecture, rather than arising from a small number of region-specific outliers. Mechanistically, this layered model is consistent with known PAR-clock coupling, in which PARP1 activity can oscillate and PARP1 can PARylate CLOCK in a time-dependent manner, linking PAR signaling to circadian state.^7^

Our genetic analyses further strengthen the biological relevance of this framework. Across ACC-relevant GWAS umbrella phenotypes spanning cognition, neurodegeneration, and sleep/circadian traits, E3 ligases mapped to disease-associated loci are disproportionately PAR-binding (**Fig. 3A-B).** A concrete precedent for how PAR binding can act as a functional “switch” for ubiquitin control is RNF146, also known as Iduna, where PAR binding activates E3 ligase function and enables PAR-dependent ubiquitination of PAR-associated targets^4^, with structural work showing WWE-domain recognition of PAR units (iso-ADP-ribose) and ligand-driven conformational activation.^34,35^

Circadian phase organization provides an additional and critical dimension to this model. PAR-binding E3 ligases occupy structured circadian windows that differ from the broader E3 background (**Fig. 4A-D).** Notably, the PAR-binding subset is enriched in transition regions of the circadian cycle rather than being confined to extreme activation or repression phases. These transition windows plausibly correspond to periods of heightened regulatory handoff—when transcriptional programs switch dominance and when protein turnover and remodeling can have an outsized impact. This interpretation aligns with the broader principle that timed ubiquitin-mediated turnover is central to clock dynamics, including well-characterized E3 control nodes such as SCF(FBXL3)/CRY stability,^13^ FBXL21 counter-regulation of CRY turnover,^36^ β-TrCP-mediated PER2 degradation,^15^ and MDM2-dependent regulation of PER2 stability and period control.^16^ Beyond these protein-stability nodes, the mammalian core clock also operates as a genome-scale transcriptional regulator, with rhythmic clock-factor occupancy linked to daily cycles in transcriptional engagement and chromatin state at many loci.^37^ Thus, if PAR-binding E3 ligases modulate core clock components in a phase-dependent way, they might indirectly influence broader chromatin and downstream transcriptional programs, though we do not measure that layer here. While our analyses are correlational, the consistency of the timing structure across representations supports the idea that PAR-binding status is linked to circadian organization rather than reflecting random rhythmicity.

Cell-type–resolved analyses further refine this picture by identifying microglia as a dominant compartment for circadian-linked and PAR-binding circadian E3 enrichment (**Fig. 5B-C).** Microglia are uniquely positioned at the intersection of immune surveillance, synaptic remodeling, and metabolic responsiveness. Beyond general “neuroinflammation” framing, microglia have direct precedent for circadian-relevant circuit remodeling via synaptic engulfment/pruning^38^ and for disease-relevant synapse loss mechanisms in Alzheimer’s models involving complement–microglia pathways.^39^ In this context, the disproportionate weighting of PAR-binding circadian E3 ligases within the microglial E3 program suggests that PAR signaling may intersect with microglial circadian state to bias ubiquitin-dependent regulation of inflammatory signaling and clearance pathways.

Several limitations of this study warrant explicit consideration. First, all analyses rely on transcriptomic proxies and curated annotations rather than direct measurements of PAR levels, ubiquitin activity, or protein turnover; expression does not equate to enzymatic activity, and circadian rhythmicity at the mRNA level may not fully reflect post-translational regulation.

Second, circadian phase annotations are derived from heterogeneous experimental systems with varying entrainment conditions, tissue sources, and rhythm-detection thresholds. More fundamentally, comprehensive time-of-day-resolved datasets for the human brain, particularly those capturing PAR synthesis, PAR–protein interactions, and ubiquitin flux, are currently unavailable. This limitation necessitates emphasis on relative phase structure and class-level organization rather than precise temporal alignment or causal inference. Despite these limitations, the framework developed here generates clear, testable predictions. (1) PAR levels and PAR-binding E3 ligase activity should exhibit coordinated circadian modulation in neural tissues, with maximal coupling during transition phases of the clock (conceptually consistent with PARP1’s circadian/metabolic coupling.^7^ (2) Perturbation of PAR synthesis or degradation should have phase-dependent effects on ubiquitination and protein stability for substrates regulated by PAR-binding ligases (with RNF146 providing a direct PAR-activated E3 precedent).^4,35^ (3) microglia-specific manipulation of PAR-binding circadian E3 ligases should preferentially impact time-of-day-dependent inflammatory and clearance phenotypes (microglia synaptic/clearance precedent.^38,39^ Lastly, (4) genetic variants that disrupt PAR–E3 interactions may sensitize neural systems to circadian misalignment, providing a mechanistic link between sleep disruption and neurodegenerative risk.

In summary, our results support a model in which non-covalent PAR-binding E3 ubiquitin ligases form a brain-deployed, circadian-structured regulatory layer that can couple PAR signaling to clock-relevant ubiquitination programs. By integrating brain expression architecture, ACC-focused human genetic convergence, circadian phase organization, and cell-type weighting, we define a prioritized set of PAR-responsive ligases positioned to influence circadian regulation through selective control of protein turnover and signaling state. A key implication is that the next mechanistic question is no longer whether PAR and circadian disruption co-occur in disease, but which PAR-binding E3 ligases directly engage clock components or clock-adjacent regulators in a PAR-dependent manner, and whether that engagement is phase- and cell-context–specific. Answering this would convert the present framework into an experimentally testable interaction map (PAR → E3 → circadian targets) and, ultimately, into translational leverage: identifying and modulating PAR-dependent E3–circadian target interfaces could provide a rational strategy to stabilize circadian outputs in disorders where circadian disruption is clinically meaningful, including neurodegenerative disease contexts.

## Author Contributions

S.M. and S.K. designed the study and wrote the paper. S.M. and S.K. collected data and performed data analysis. S.M. and S.K. performed statistic review. S.M. and S.K. designed and modified the data visualization. S.K. supervised the project.

## Supporting information

All Supplemental Tables

## Acknowledgement

S.M. was supported by Northwestern Medicine Central DuPage Hospital and Delnor Hospital Medical Staff Scholarship. S.U.K. was supported by the National Institutes of Health (NIH)/National Institute of Neurological Disorders and Stroke (NINDS) R01 NS123456.

## Data Availability Statement

The original contributions presented in this study are included in the article/supplementary material. Further inquiries can be directed to the corresponding author(s).

